# *De novo* genome assembly of the meadow brown butterfly, *Maniola jurtina*

**DOI:** 10.1101/715243

**Authors:** Kumar Saurabh Singh, David J. Hosken, Nina Wedell, Richard ffrench-Constant, Chris Bass, Simon Baxter, Konrad Paszkiewicz, Manmohan D Sharma

## Abstract

**Background:** Meadow brown butterflies (*Maniola jurtina*) on the Isles of Scilly represent an ideal model in which to dissect the links between genotype, phenotype and long-term patterns of selection in the wild - a largely unfulfilled but fundamental aim of modern biology. To meet this aim, a clear description of genotype is required.

**Findings:** Here we present the draft genome sequence of *M. jurtina* to serve as an initial genetic resource for this species. Seven libraries were constructed using DNA from multiple wild caught females and sequenced using Illumina, PacBio RSII and MinION technology. A novel hybrid assembly approach was employed to generate a final assembly with an N50 of 214 kb (longest scaffold 2.9 Mb). The genome encodes a total of 36,294 genes. 90.3% and 88.7% of core BUSCO (Benchmarking Universal Single-Copy Orthologs) Arthropoda and Insecta gene sets were recovered as complete single-copies from this assembly. Comparisons with 17 other Lepidopteran species placed 86.5% of the assembled genes in orthogroups.

**Conclusions:** Our results provide the first high-quality draft genome and annotation of the butterfly *M. jurtina.*

## Background Information

The Meadow brown butterfly (*Maniola jurtina*, NCBI:txid191418) is a member of the nymphalid subtribe satyrini. It is an important model organism for the study of lepidopteran ecology, evolution and has been extensively studied by ecological geneticists for many years [1-3]. Found across the Palearctic realm it primarily habituates in grasslands, woodland rides, field-margins and can even be found in overgrown gardens.

The species displays marked sexual dimorphism. Females are more colourful than males and have large upper-wing eyespots (Figure 1). It also exhibits considerable quantitative variation in the sub-marginal spot pattern of its wings [4] and therefore represents an ideal model in which to dissect the links between genotype, phenotype and long-term patterns of selection in the wild [5] - a largely unfulfilled but fundamental aim of modern biology. This draft genome and corresponding annotations will offer a core resource for ongoing work in lepidopterans and other arthropods of ecological importance.

**Figure 1:**
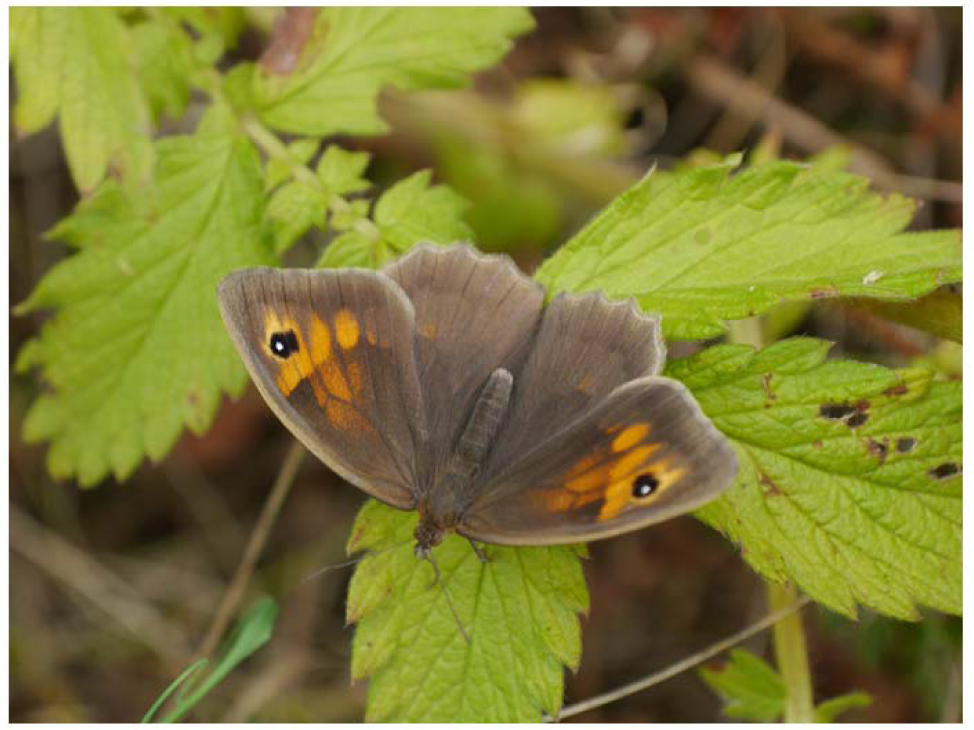
Female *Maniola jurtina* (picture credit: Richard ffrench-Constant)

## Data Description

### Sampling and sequencing

Adult meadow brown (*Maniola jurtina*) butterflies were collected from the wild (Isles of Scilly, Cornwall) in June 2012, anaesthetized by refrigeration for 2 hours and then killed by subsequent freezing. High molecular weight genomic DNA was extracted from whole body (excluding wings) of five individual females using the genomic-tip 100/G kit (Qiagen, Hilden, Germany) supplemented with RNase A (Qiagen, Hilden, Germany) and Proteinase K (New England Biolabs, Hitchin, UK) treatment, as per the manufacturer’s instructions. DNA quantity and quality were subsequently assessed using a NanoDrop-2000 (Thermo Scientific™, Loughborough, UK) and a Qubit 2.0 fluorometer (Life Technologies). Molecular integrity was assessed using pulse-field gel electrophoresis.

Illumina data (100bp paired-end) was generated using standard Illumina protocols for a 250-500 bp PE library and 3-5 KB, 5-7 KB mate-pair libraries (Table S1). 20 kb PacBio libraries were generated and size-selected following the manufacturers recommended protocols and sequenced on 18 SMRT cells of the RSII instrument. Finally, exceptionally long reads (longest read 300Kb) were obtained using the Oxford Nanopore Technologies MinION platform (R7.4) (Table S2). Illumina, PacBio and MinION library preparation and sequencing were performed by the Exeter Sequencing Service, University of Exeter.

### Genome characteristics

The genome characteristics of *M. jurtina* were estimated using a k-mer based approach implemented in GenomeScope [5]. Short-read Illumina reads were quality filtered and subjected to 19-mer frequency distribution analysis using Jellyfish -*v*2.2.0 [6]. The 19-mer profile showed a unimodal distribution of k-mers with a coverage peak at 17. The estimated genome size was approximately 574 Mbp. The estimated heterozygosity rate was 2% and the genome is comprised of 75.61% repetitive elements that are likely to contain units of highly repetitive W chromosome as the samples used in this study were all female (Table S3). Note here that although the genome size estimated via this method is strongly dependent on the coverage profile, based on the genome size of the most closely related species *Bicyclus anynana* (475 Mb), this estimate does not seem inordinate.

### Genome assembly

Genome assembly was performed by adopting a novel hybrid approach (Figure 2). Paired-end Illumina reads were trimmed and filtered for quality values using Trim Galore *-v*0.4.2 [7] and assembled using Spades *-v*3.9.1 [8] and Redundans *-v*0.14c [9]. Long reads obtained from MinION were mixed with PacBio reads and assembled using Canu *-v*1.3.0 [10]. The short and long read assemblies were later merged using QuickMerge *-v*0.3.0 [11] and redundancy reduction, scaffolding and gap closing was carried using Redundans. The spades diploid assembly was used as input together with Illumina paired-end and mate-pair data as required by Redundans for scaffolding. The draft assembly was polished using arrow (part of the genomicconsensus package in PacBio tools) -*v*2.3.2 [12], which exclusively mapped long PacBio reads against the draft assembly using the BLASR pipeline [13]. The draft assembly was also polished with the Illumina short-reads using Pilon *-v*1.23.0 [14]. A summary of the different assembly steps is available in Table 2 and the final assembly properties are detailed in Table S4.

**Figure 2:**
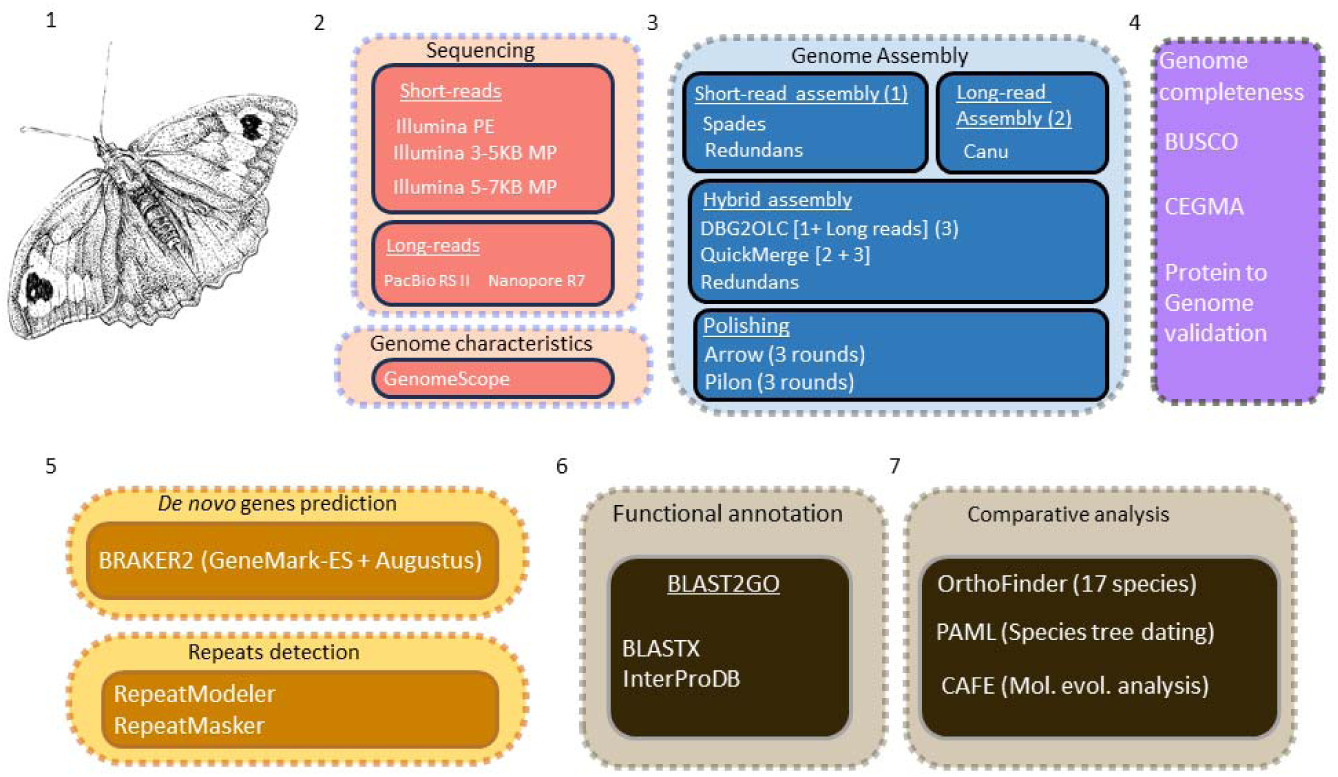
Schematic overview of the workflow used for sequencing, genome size estimation, assembly and annotation of the *M. jurtina* genome. Transcriptome data (orange segment) were obtained from publicly available sources at NCBI and only used for genome annotation.

**Table 1.**
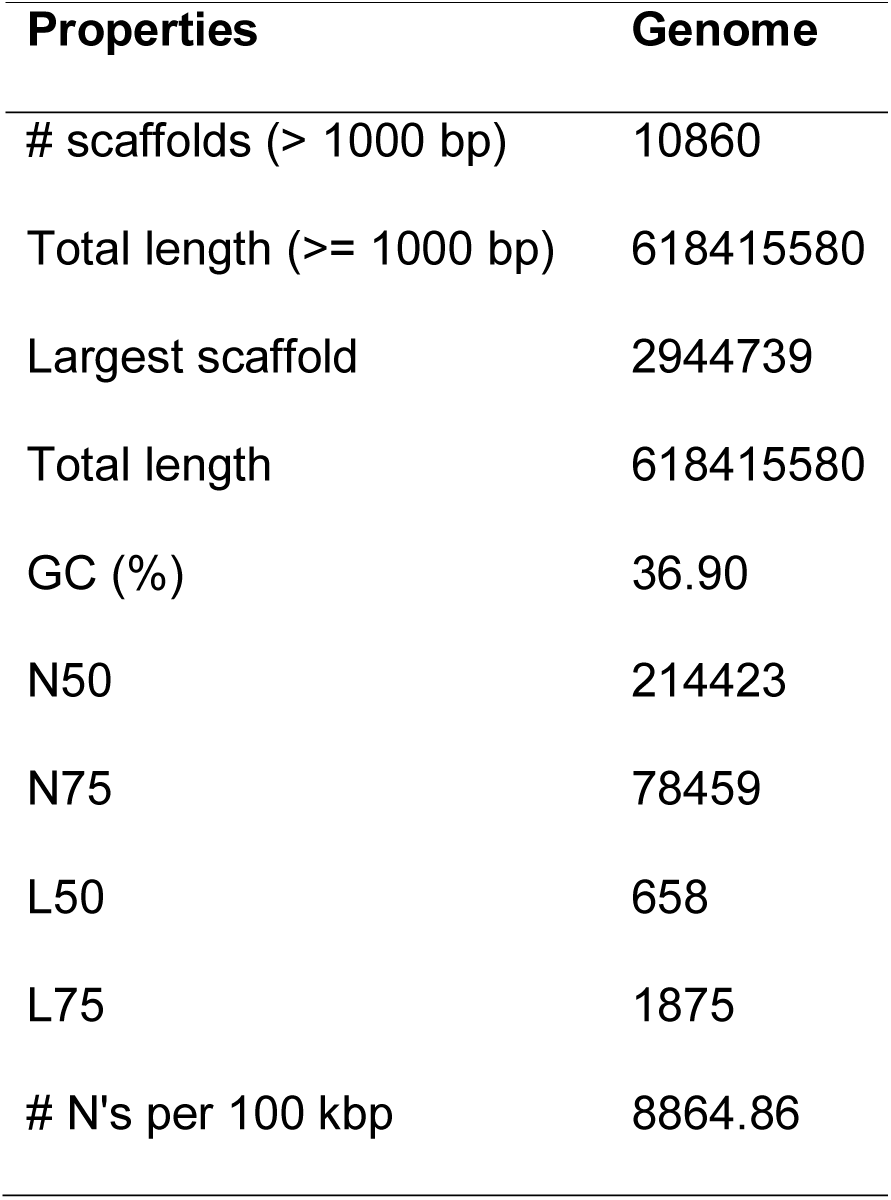
*M. jurtina* genome properties

**Table 2.**
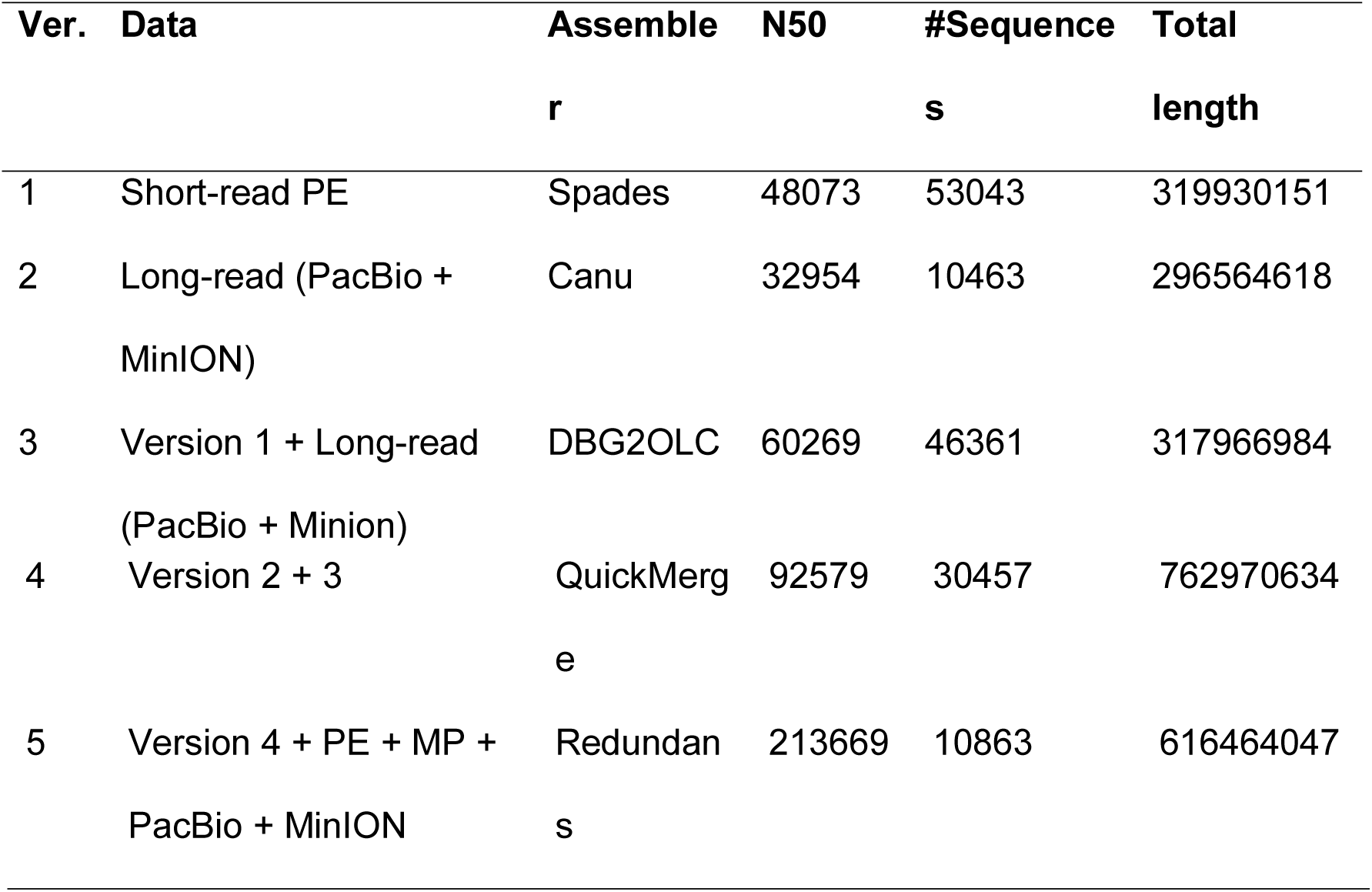
Different assembly versions (ver), data, software used and summary statistics

### Evaluation of the completeness of the genome assembly

The completeness of the draft genome was assessed by mapping raw short and long reads against the assembly. The percentage of aligned reads ranged from 94 to 95% using paired-end and mate-paired short reads. BUSCO (Benchmarking universal single-copy orthologs) *-v*3.0.2 [15] and CEGMA (Core Eukaryotic genes mapping approach) *-v*2.5.0 [16] were used to check genomic completeness of the assembly. In the case of BUSCO, Arthropoda and Insecta gene sets were compared against the assembly. About 91.5% and 88% total BUSCO genes were identified in the Arthropoda and Insecta sets respectively. Additionally, 91% CEGMA genes, both complete and partials, were successfully found in the assembly (Table S5 and S6). We also assessed the completeness of this assembly by comparing the *M. jurtina* genome against *H. melpomene* and *B. anynana* (a close relative). Number of genomic sequence matches found between *M. jurtina* and *B. anynana* were significantly more as compared to *H. melpomene*. Number of matches found in this comparison correlates well with genome size and species divergence times. The genome size of *H. melpomene* (∼250 MB) is relatively smaller than *B. anynana* (∼475 MB). Therefore many *M. jurtina* genomic sequences ended up with no hits.

### Genome Annotation

Before predicting gene models, the genome of *M. jurtina* was masked for repetitive elements using RepeatMasker -*v*4.0.7 [17]. RepeatModeler *-v*1.0.11 [18] was used to model the repeat motifs and transposable elements. Repeats originating from coding regions were removed by performing a BLAST search against the *B. anynana* proteins. Sequence with hits at E-value > 1e^−10^ were filtered out. The RepBase *-v*24.05 library was then merged with the repeats predicted by RepeatModeler and used to mask the *M. jurtina* genome. Protein coding genes were predicted using GeneMark-ES *-v*4.3.8 [19] and AUGUSTUS -*v*3.3.0 [20] implemented in the BRAKER *-v* 2.1.2 [21] pipeline using RNA-seq alignments as evidence. Publicly available *M. jurtina* RNA-seq datasets (SRR3724201, SRR3724266, SRR3724269, SRR3724271, SRR3724198, SRR3724196, SRR3724195, SRR3721773, SRR3721752, SRR3721684, SRR3721695) were downloaded from NCBI and mapped individually against the repeat masked genome using STAR -*v*2.7.1 [22]. The bam files from individual samples were then combined and fed into BRAKER. Low quality genes with fewer than 50 amino-acid and/or exhibiting premature termination were removed from the gene set yielding a final count of 38101 genes. In the final gene set, mean gene length, mean CDS length, mean intron length and exon number per protein were 4144 bp, 976 bp, 921 bp and 5 respectively (Table S7).

Functional annotation of the *de-novo* predicted gene models was carried out using homology searches against the NCBI nr database and Interpro database using BLAST2GO *-v*5.2.5 [23]. Approximately 34263 out of 38101 genes (90%) of the predicted genes could be assigned functional annotation based on BLAST searches against the non-redundant protein database of NCBI and Interpro.

### Comparison to other Lepidopteran species

To characterize orthology and investigate gene family evolution across Lepidoptera, the final annotation set for *M. jurtina* was compared to 17 other genomes including a dipteran (*Drosophila melanogaster*), and a trichopteran (*Limnephilus lunatus*) as outgroups. The proteomes of *Amyelois transitella* v1.0, *B. anynana* v1.2, *Bombyx mori* v1.0, *Calycopis cecrops* v1.1, *Chilo suppressalis* v1.0, *Danaus plexippus* v3.0, *Heliconius melpomene* v2.0, *Junonia coenia* v1.0, *Limnephilus lunatus* v1.0, *Melitaea cinxia, Operophtera brumata* v1.0, *Papilio polytes* v1.0, *Phoebis sennae* v1.1, *Plodia interpunctella* v1.0, *Plutella xylostella* v1.0 were downloaded from Lepbase. OrthoFinder *-v*1.1.8 [24] was used to define orthologous groups (gene families) of genes between these peptide sets. 50% of all genes were in orthogroups with 23 or more genes and were contained in the largest 4439 orthogroups. The combined gene count of these species was 349442 of which 86.5% were assigned to 15064 orthogroups. There were 2915 orthogroups with all species present and 39 of these consisted entirely of single-copy genes. A total of 216 gene families were specific to *M. jurtina* compared to 627 and 1716 in butterfly and moths respectively (Figure 3A).

**Figure 3:**
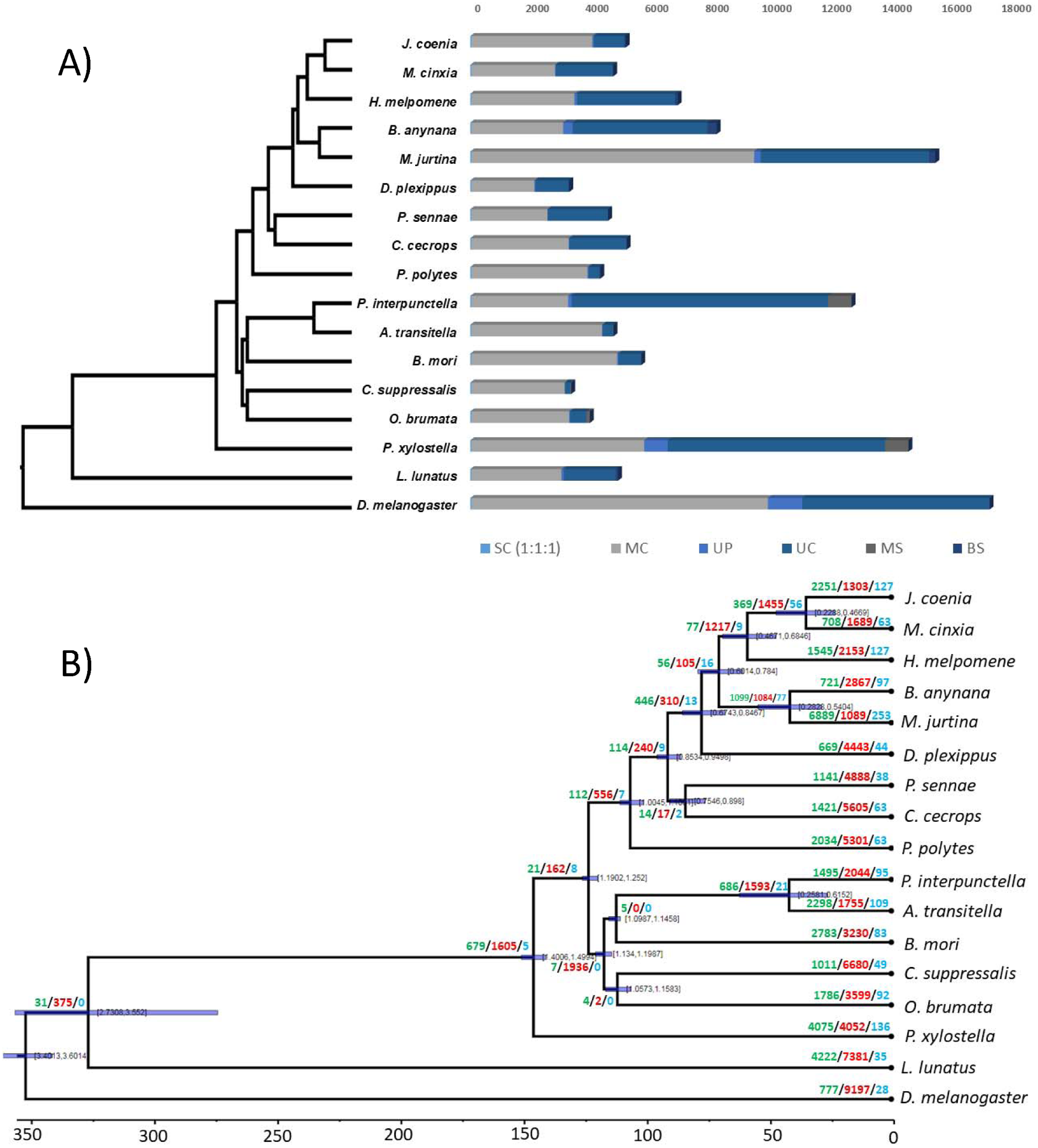
Evolutionary and comparative genomic analysis. **(A)** Ortholog analysis of *M. jurtina* with 16 other arthropod species. SC indicates common orthologs with the same number of copies in different species, MC indicates common orthologs with different copy numbers in different species, UP indicates species specific paralogs, UC indicates all genes which were not assigned to a gene family, MS indicates moths specific genes and BS indicates butterfly specific genes. **(B)** Species phylogenetic tree and gene family evolution. Numbers on the node indicate counts of the gene families that are expanding (green), contracting (red) and rapidly evolving (blue).

### Phylogenetic tree construction and divergence time estimation

Phylogenetic analysis was performed using 39 single-copy orthologous genes from common gene families found by OrthoFinder (Table S8). Additionally, OrthoFinder generated a species tree where *D. melanogaster* was used as the outgroup. The species tree was rooted using the STRIDE -*v*1.0.0 [25] algorithm implemented in OrthoFinder. MCMCTREE, as implemented in PAML -v4.9e [26], was then used to estimate the divergence times of *M. jurtina* with approximate likelihood calculation. For this, the substitution rate was estimated using *codeml* by applying root divergence age between the Diptera, Lepidoptera and Trichoptera as 350 MY [27]. This is a simple fossil calibration of 350 MY for the root. The estimated substitution rate was the per site substitution rate for the amino acid data set and used to set priors for the mean substitution rate in Bayesian analysis. As a second step, the gradient (g) and Hessian (H) of branch lengths for all 17 species were also estimated. Finally, the tree file with fossil calibrations (Table S9), the gradient vector and hessian matrices file and the concatenated genes alignment information were used in the approximate likelihood calculation. The parameter settings of MCMCTREE were as follows: clock = 2, model = 3, BDparas = 110, kappa_gamma = 6 2, alpha_gamma = 11, rgene_gamma = 9.09, and sigma2_gamma = 1 4.5. The phylogenetic analysis showed that *M. jurtina* is more closely related to *B. anynana* than to *H. melpomene* or *M. cinxia.* The estimated divergence time between *M. jurtina* and *B. anynana* was estimated to be around 34 MYA and that between *M. jurtina* and *H. melpomene* is estimated as 57 MYA (Figure 3B). Whole genome alignments, using Mummer *-v*3.1.0 [28] between *M. jurtina* – *B. anynana* and *M. jurtina* - *H. melpomene* were also performed to confirm this relatedness (Figure 4). In the dated phylogeny, the most species rich family Nymphalidae has remained stable and diverged from Papilionidae around 90 MY ago. This age is also supported by previously published butterfly phylogenies [29, 30].

**Figure 4:**
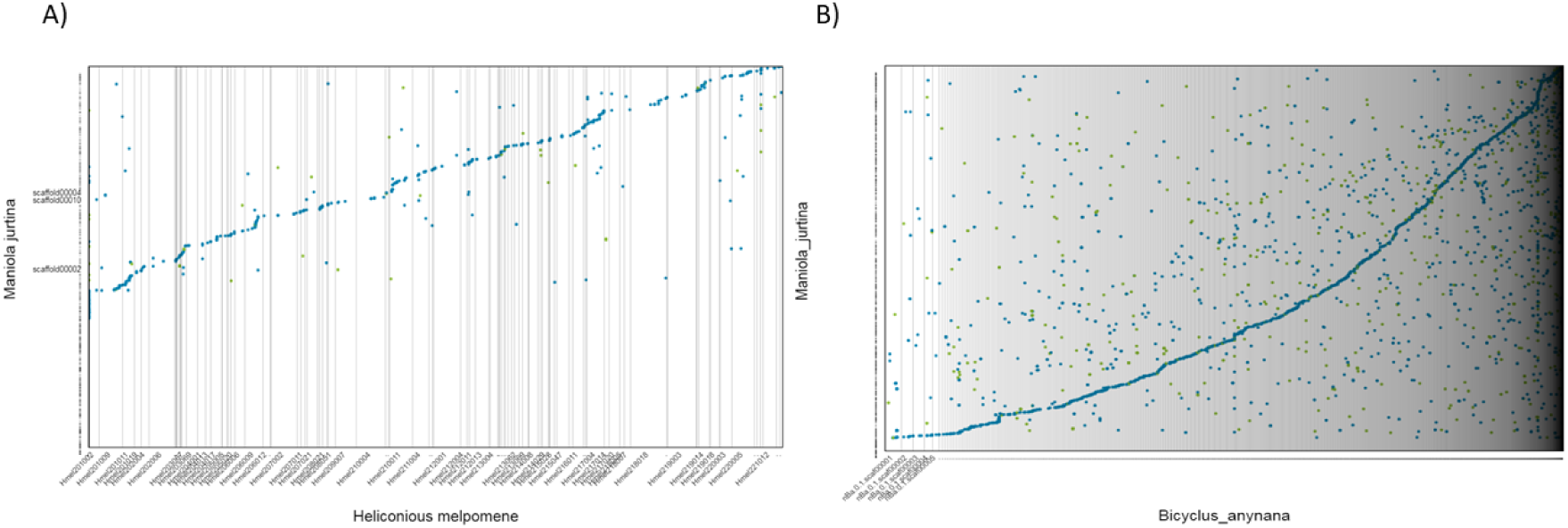
Genome comparisons. *Comparison of* the *Maniola jurtina genome with Heliconious melpomene and Bicyclus anynana*. The dot plots were generated using Mummer. The plots show relatedness of *M. jurtina* with A) *H. melpomene* and B) *B. anynana*. Both of these genomes were taken as references (x-axis) and queried using *M. jurtina* (y-axis) genome. Plot B shows more consistent and contiguous alignments than plot A. The dot plots were generated using https://dnanexus.github.io/dot/

### Analysis of gene family evolution

Gene family evolution across arthropods was investigated using CAFE *-v*3.0 [31]. CAFE models the evolution of gene family size across a species phylogeny under a ML birth–death model of gene gain and loss and simultaneously reconstructs ML ancestral gene family sizes for all internal nodes, allowing the detection of expanded gene families within lineages. We ran CAFE on our matrix of gene family sizes generated by OrthoFinder and modelled their evolution along the dated species tree. Genes involved in binding, metabolism and transport of natural or synthetic allelochemicals are particularly found to be rapidly evolving in *M. jurtina* (Figure 3B).

### Analysis of spot pattern related genes

Dowdeswell, Fisher and Ford first studied the island-specific wing-spot patterns in *M. jurtina* on the isles of Scilly [3], and this work was continued for more than 20 years (reviewed in [1]). Their major findings, which became a cornerstone of ecological genetics, have been re-visited and largely re-confirmed with contemporary data [32]. Patterns of wing-spot polymorphism have remained unchanged on some islands over 60 years and there is some evidence of genetic differentiation across the Scillies [32]. Nonetheless, much remains to be done to better understand the underlying genetics of spot pattern variation in this species.

Butterfly wing patterns have long been suggested to be polygenic [33] and recent evidence from *B. anynana* (very closely related to *M. jurtina*) has confirmed this to be the case and strongly suggested that 10-11 different genomic regions may be involved in eye-spot number variation [34] and see [35].

To test whether any genes involved in wing or spot -pattern formation across Lepidoptera were identifiable in the current *Maniola* assembly, we first performed a wide literature search PubMed (https://www.ncbi.nlm.nih.gov/pubmed/) using the keywords *Lepidoptera, butterfly, wing, spot, pattern, gene* and then manually filtered through the results to generate a list of candidate genes (Table 3).

**Table 3.** Candidate wing spot patterning genes obtained from a literature search are listed in column 1. Column 2 has the number of annotated orthologues across Lepidoptera in our NCBI protein database search using the full gene name as listed and alternate names (comma separated). Column 3 presents the number of proteins per gene that matched in our Exonerate workflow.

|  | Candidate gene<br>(*alt. name in brackets*) [Reference] | NCBI Protein<br>Matches within<br>Lepidoptera (*n*) | protein2genome<br>matches against<br>the *Maniola*<br>genome (*n*) |
| --- | --- | --- | --- |
| 1 | *Ap* [37] | 1 | 0 |
| 2 | *APC* [38] | 18 | 4 |
| 3 | *Apterous* / *apterousA* (*apA*) [37, 39] | 25 | 4 |
| 4 | *Black* [40] | 5 | 0 |
| 5 | *C2 domain-containing protein 5* (*C2CD5*) [34] | 32 | 7 |
| 6 | *Calcium-activated potassium channel slowpoke*<br>(*slo*) [34, 41] | 205 | 191 |
| 7 | *Cortex* [42, 43] | 26 | 0 |
| 8 | *Decapentaplegic (dpp)* [44, 45] | 145 | 86 |
| 9 | *Distal-less (Dll)* [44, 46, 47] | 47 | 1 |
| 10 | *Doublesex (dsx)* [48, 49] | 204 | 101 |
| 11 | *DP transcription factor (dp)* [37] | 4 | 2 |
| 12 | *Ecdysone receptor (ecr)* [46, 50] | 101 | 33 |
| 13 | *engrailed (en)* [51] | 19 | 0 |
| 14 | *Geranylgeranyl pyrophosphate synthase (GGPS1)* [34] | 26 | 10 |
| 15 | *Hox* [52] | 39 | 6 |
| 16 | *Invected* [51] | 15 | 0 |
| 17 | *Naked cuticle* [38] | 16 | 3 |
| 18 | *Neutral Ceramidase (Cdase)* [34, 41] | 32 | 22 |
| 19 | *Notch (N)* [47] | 117 | 72 |
| 20 | *numb* [34] | 40 | 0 |
| 21 | *Optix* [42, 53] | 5 | 3 |
| 22 | *Phosphoinositide 3-kinase adapter protein 1 (PIK3AP1)* [34] | 58 | 0 |
| 23 | *poikilomousa / poik (HM00025)* [54] | 2 | 0 |
| 24 | *spaetzle (spz)* [34, 41] | 65 | 2 |
| 25 | *spalt (Sal)* [51] | 1 | 1 |
| 26 | *Transient receptor potential channel pyrexia (pyx)* [34, 41] | 74 | 23 |
| 27 | *Vermilion* [37] | 2 | 0 |
| 28 | *wingless (wg)* [55] | 0 | - |
| 29 | *WntA* [42] | 1 | 1 |
| 30 | *Zinc finger CCCH domain-containing protein 10*<br>(ZC3H10) [34] | 23 | 1 |

This includes a selection of regulators possibly responsible for pattern variation (*APC, Naked cuticle*), transcription factors linked with eyespot patterning (*Distal-less, Dll*, and *Engrailed, En*), along with other transcription regulators such as *Apterous* and *DP*. Additionally, we considered *poik* (HM00025), also known as *cortex, Optix, Doublesex, Hox, Vermilion* and *black* (pigment synthesis) along with the *Ecdysone receptor* (*EcR*) involved in wing pattern plasticity.

NCBI esearch and efetch tools were used to filter (*NOT partial NOT hypothetical NOT uncharacterized*), and query individual spot pattern proteins across Lepidoptera using both, full and abbreviated protein names where available (Table 3; total 1347 homologues) and then these proteins were queried against the *Maniola* genome using Exonerate -v2.2.0 [36] protein2genome model with the following customised options *--refine region --score 900 --percent 70 -S FALSE --softmasktarget TRUE -- bestn 1 --ryo \*”>%*ti (*%*tab* - %*tae) coding (*%*tcb* - %*tce) cds_length (*%*tcl)\n*%*tcs\n*”.

Protein to genome matches were found for 20 out of the 30 candidate genes (Table 3). We further cross checked this by creating a blast database of the 1347 homologue spot pattern related proteins from Lepidoptera and then searching the homologues within the *M. jurtina* proteome for matches. This resulted in over 1500 matches (see supplementary file 2).

Specific experiments now need to be undertaken to further test candidate genes and their possible roles in wing-spot polymorphism, and to revisit other findings from Ford and co-workers (reviewed in [1]) in the iconic Scillies study system.

## Conclusion

Here we present a high-quality draft assembly and annotation of the butterfly *M. jurtina.* The assembly, along with the cross-species comparisons and elements of key spot-pattern genes will offer a core genomic resource for ongoing work in lepidopterans and other arthropods of ecological importance.

## Availability of Data and Materials

The raw sequencing data and genome assembly have been deposited at the NCBI SRA database under the BioProject PRJNA498046 and genome accession number VMKL00000000. Blast results, annotation and proteome associated with this manuscript are available at https://zenodo.org/record/3352197.

## Acknowledgements

The authors would like to acknowledge the use of University of Exeter’s Advanced Computing Resources and Maisy Inston (University of Exeter) for the graphical illustration of a *M. jurtina* sample. There are no competing interests to declare.

## References

1. Ford EB. Ecological genetics. London: Methuen Ltd.; 1965.

2. Dowdeswell W. Experimental studies on natural selection in the butterfly, Maniola jurtina. Heredity. 1961;16 1:39.

3. Dowdeswell WH, Fisher RW and Ford EB. The quantitative study of populations in the Lepidoptera; *Maniola jurtina L*. Heredity (Edinb). 1949;3 Pt. 1:67–84. doi:10.1038/hdy.1949.3.

4. Brakefield PM and van Noordwijk AJ. The Genetics of Spot Pattern Characters in the Meadow Brown Butterfly *Maniola jurtina* (Lepidoptera, Satyrinae). Heredity. 1985;54 Apr:275–84. doi:DOI 10.1038/hdy.1985.37.

5. Baxter SW, Hoffman JI, Tregenza T, Wedell N and Hosken DJ. EB Ford revisited: assessing the long-term stability of wing-spot patterns and population genetic structure of the meadow brown butterfly on the Isles of Scilly. Heredity. 2017;118 4:322–9. doi:10.1038/hdy.2016.94.

6. Vurture GW, Sedlazeck FJ, Nattestad M, Underwood CJ, Fang H, Gurtowski J, et al. GenomeScope: fast reference-free genome profiling from short reads. Bioinformatics. 2017;33 14:2202–4. doi:10.1093/bioinformatics/btx153.

7. Marcais G and Kingsford C. A fast, lock-free approach for efficient parallel counting of occurrences of k-mers. Bioinformatics. 2011;27 6:764–70. doi:10.1093/bioinformatics/btr011.

8. Krueger F. TrimGalore; https://github.com/FelixKrueger/TrimGalore. xThe Babraham Institute, 2016.

9. Bankevich A, Nurk S, Antipov D, Gurevich AA, Dvorkin M, Kulikov AS, et al. SPAdes: A New Genome Assembly Algorithm and Its Applications to Single-Cell Sequencing. J Comput Biol. 2012;19 5:455–77. doi:10.1089/cmb.2012.0021.

10. Pryszcz LP and Gabaldon T. Redundans: an assembly pipeline for highly heterozygous genomes. Nucleic Acids Res. 2016;44 12:e113. doi:10.1093/nar/gkw294.

11. Koren S, Walenz BP, Berlin K, Miller JR, Bergman NH and Phillippy AM. Canu: scalable and accurate long-read assembly via adaptive k-mer weighting and repeat separation. Genome Res. 2017;27 5:722–36. doi:10.1101/gr.215087.116.

12. Chakraborty M, Baldwin-Brown JG, Long AD and Emerson JJ. Contiguous and accurate de novo assembly of metazoan genomes with modest long read coverage. Nucleic Acids Res. 2016;44 19:e147. doi:10.1093/nar/gkw654.

13. Pacific Biosciences of California I. GenimicConsensus; PacBio® tools-https://github.com/PacificBiosciences/GenomicConsensus.

14. Chaisson MJ and Tesler G. Mapping single molecule sequencing reads using basic local alignment with successive refinement (BLASR): application and theory. BMC Bioinformatics. 2012;13 1:238. doi:10.1186/1471-2105-13-238.

15. Walker BJ, Abeel T, Shea T, Priest M, Abouelliel A, Sakthikumar S, et al. Pilon: an integrated tool for comprehensive microbial variant detection and genome assembly improvement. PLoS One. 2014;9 11:e112963. doi:10.1371/journal.pone.0112963.

16. Simao FA, Waterhouse RM, Ioannidis P, Kriventseva EV and Zdobnov EM. BUSCO: assessing genome assembly and annotation completeness with single-copy orthologs. Bioinformatics. 2015;31 19:3210–2. doi:10.1093/bioinformatics/btv351.

17. Parra G, Bradnam K and Korf I. CEGMA: a pipeline to accurately annotate core genes in eukaryotic genomes. Bioinformatics. 2007;23 9:1061–7. doi:10.1093/bioinformatics/btm071.

18. Smit A, Hubley, R & Green, P. RepeatMasker Open-4.0. http://www.repeatmasker.org. 2013-2015.

19. Smit A, Hubley, R. RepeatModeler Open-1.0. http://www.repeatmasker.org. 2008-2015.

20. Lomsadze A, Ter-Hovhannisyan V, Chernoff YO and Borodovsky M. Gene identification in novel eukaryotic genomes by self-training algorithm. Nucleic Acids Res. 2005;33 20:6494-506. doi:10.1093/nar/gki937.

21. Stanke M and Morgenstern B. AUGUSTUS: a web server for gene prediction in eukaryotes that allows user-defined constraints. Nucleic Acids Research. 2005;33:W465-W7. doi:10.1093/nar/gki458.

22. Hoff KJ, Lange S, Lomsadze A, Borodovsky M and Stanke M. BRAKER1: Unsupervised RNA-Seq-Based Genome Annotation with GeneMark-ET and AUGUSTUS. Bioinformatics. 2016;32 5:767-9. doi:10.1093/bioinformatics/btv661.

23. Dobin A, Davis CA, Schlesinger F, Drenkow J, Zaleski C, Jha S, et al. STAR: ultrafast universal RNA-seq aligner. Bioinformatics. 2013;29 1:15–21. doi:10.1093/bioinformatics/bts635.

24. Gotz S, Garcia-Gomez JM, Terol J, Williams TD, Nagaraj SH, Nueda MJ, et al. High-throughput functional annotation and data mining with the Blast2GO suite. Nucleic Acids Research. 2008;36 10:3420–35. doi:10.1093/nar/gkn176.

25. D.M. E and S. K. OrthoFinder2: fast and accurate phylogenomic orthology analysis from gene sequences. bioRxiv. 2018:466201. doi:10.1101/466201.

26. Emms DM and Kelly S. STRIDE: Species Tree Root Inference from Gene Duplication Events. Molecular biology and evolution. 2017;34 12:3267-78. doi:10.1093/molbev/msx259.

27. Yang Z. PAML 4: phylogenetic analysis by maximum likelihood. Molecular biology and evolution. 2007;24 8:1586-91. doi:10.1093/molbev/msm088.

28. Kjer KM, Ware JL, Rust J, Wappler T, Lanfear R, Jermiin LS, et al. Response to Comment on “Phylogenomics resolves the timing and pattern of insect evolution”. Science. 2015;349 6247:487-. doi:10.1126/science.aaa7136.

29. Kurtz S, Phillippy A, Delcher AL, Smoot M, Shumway M, Antonescu C, et al. Versatile and open software for comparing large genomes. Genome Biol. 2004;5 2 doi:ARTN R12; DOI 10.1186/gb-2004-5-2-r12.

30. Wahlberg N, Wheat CW and Pena C. Timing and Patterns in the Taxonomic Diversification of Lepidoptera (Butterflies and Moths). Plos One. 2013;8 11 doi:ARTN e80875; 10.1371/journal.pone.0080875.

31. Espeland M, Breinholt J, Willmott KR, Warren AD, Vila R, Toussaint EFA, et al. A Comprehensive and Dated Phylogenomic Analysis of Butterflies. Curr Biol. 2018;28 5:770-+. doi:10.1016/j.cub.2018.01.061.

32. De Bie T, Cristianini N, Demuth JP and Hahn MW. CAFE: a computational tool for the study of gene family evolution. Bioinformatics. 2006;22 10:1269–71. doi:10.1093/bioinformatics/btl097.

33. Beldade P and Brakefield PM. The genetics and evo-devo of butterfly wing patterns. Nature Reviews Genetics. 2002;3 6:442–52. doi:10.1038/nrg818.

34. Rivera-Colón AG, Westerman EL, Van Belleghem SM, Monteiro A and Papa R. The genetic basis of hindwing eyespot number variation in <em>Bicyclus anynana</em> butterflies. bioRxiv. 2018:504506. doi:10.1101/504506.

35. Monteiro A and Prudic KM. Multiple approaches to study color pattern evolution in butterflies. Trends in Evolutionary Biology. 2010;2 1:e2-e.

36. Slater GS and Birney E. Automated generation of heuristics for biological sequence comparison. BMC Bioinformatics. 2005;6:31. doi:10.1186/1471-2105-6-31.

37. Beldade P, Brakefield PM and Long AD. Generating phenotypic variation: prospects from “evo-devo” research on Bicyclus anynana wing patterns. Evol Dev. 2005;7 2:101–7. doi:10.1111/j.1525-142X.2005.05011.x.

38. Saenko SV, Brakefield PM and Beldade P. Single locus affects embryonic segment polarity and multiple aspects of an adult evolutionary novelty. Bmc Biol. 2010;8 doi:Artn 111; 10.1186/1741-7007-8-111.

39. Prakash A and Monteiro A. apterous A specifies dorsal wing patterns and sexual traits in butterflies. Proc Biol Sci. 2018;285 1873 doi:10.1098/rspb.2017.2685.

40. Walker JF and Monteiro A. Determining the putative source of a morphogen underlying black spot development in Pieris rapae butterflies. Integrative and Comparative Biology. 2013;53:E389–E.

41. Ozsu N and Monteiro A. Wound healing, calcium signaling, and other novel pathways are associated with the formation of butterfly eyespots. BMC Genomics. 2017;18 1:788. doi:10.1186/s12864-017-4175-7.

42. Callier V. How the butterfly got its spots (and why it matters). P Natl Acad Sci USA. 2018;115 7:1397–9. doi:10.1073/pnas.1722410115.

43. Nadeau NJ, Pardo-Diaz C, Whibley A, Supple MA, Saenko SV, Wallbank RW, et al. The gene cortex controls mimicry and crypsis in butterflies and moths. Nature. 2016;534 7605:106.

44. Monteiro A, Chen B, Ramos DM, Oliver JC, Tong X, Guo M, et al. Distal-less regulates eyespot patterns and melanization in Bicyclus butterflies. J Exp Zool B Mol Dev Evol. 2013;320 5:321–31. doi:10.1002/jez.b.22503.

45. Connahs H, Tlili S, van Creij J, Loo TY, Banerjee T, Saunders TE, et al. Disrupting different Distal-less exons leads to ectopic and missing eyespots accurately modeled by reaction-diffusion mechanisms. bioRxiv. 2017:183491.

46. Koch PB, Merk R, Reinhardt R and Weber P. Localization of ecdysone receptor protein during colour pattern formation in wings of the butterfly Precis coenia (Lepidoptera: Nymphalidae) and co-expression with Distal-less protein. Dev Genes Evol. 2003;212 12:571–84. doi:10.1007/s00427-002-0277-5.

47. Reed RD and Serfas MS. Butterfly wing pattern evolution is associated with changes in a Notch/Distal-less temporal pattern formation process. Curr Biol. 2004;14 13:1159–66. doi:10.1016/j.cub.2004.06.046.

48. Nishikawa H, Iijima T, Kajitani R, Yamaguchi J, Ando T, Suzuki Y, et al. A genetic mechanism for female-limited Batesian mimicry in Papilio butterfly. Nat Genet. 2015;47 4:405–U169. doi:10.1038/ng.3241.

49. Kunte K, Zhang W, Tenger-Trolander A, Palmer DH, Martin A, Reed RD, et al. doublesex is a mimicry supergene. Nature. 2014;507 7491:229-+. doi:10.1038/nature13112.

50. Koch PB, Brakefield PM and Kesbeke F. Ecdysteroids control eyespot size and wing color pattern in the polyphenic butterfly Bicyclus anynana (Lepidoptera: Satyridae). J Insect Physiol. 1996;42 3:223–30. doi:Doi 10.1016/0022-1910(95)00103-4.

51. Brunetti CR, Selegue JE, Monteiro A, French V, Brakefield PM and Carroll SB. The generation and diversification of butterfly eyespot color patterns. Curr Biol. 2001;11 20:1578–85. doi:Doi 10.1016/S0960-9822(01)00502-4.

52. Hombria JC. Butterfly eyespot serial homology: enter the Hox genes. Bmc Biol. 2011;9:26. doi:10.1186/1741-7007-9-26.

53. Reed RD, Papa R, Martin A, Hines HM, Counterman BA, Pardo-Diaz C, et al. optix drives the repeated convergent evolution of butterfly wing pattern mimicry. Science. 2011;333 6046:1137–41. doi:10.1126/science.1208227.

54. Nadeau NJ, Pardo-Diaz C, Whibley A, Supple M, Wallbank R, Wu GC, et al. The origins of a novel butterfly wing patterning gene from within a family of conserved cell cycle regulators. bioRxiv. 2015:016006. doi:10.1101/016006.

55. Monteiro A, Glaser G, Stockslager S, Glansdorp N and Ramos D. Comparative insights into questions of lepidopteran wing pattern homology. BMC Developmental Biology. 2006;6 doi:Artn 52; 10.1186/1471-213x-6-52.

