## Supplemental file 1 for "*De novo* genome assembly of the meadow brown butterfly, *Maniola jurtina*"

**Supplementary Table 1**: Short-read data summary

| **Library** | **Type** | **#raw reads** | **#raw bases (bp)** | **#processed reads** | **Mean Insert size in bp (observed)** | **GC%** |
| --- | --- | --- | --- | --- | --- | --- |
| 100 | PE | 65622134 | 6562213400 | 64452732 | 203 | 36.0 |
| 1263 | MP | 280482300 | 28048230000 | 280481030 | 711 | 37.0 |
| 3k-5k | MP | 109759518 | 10975951800 | 108327272 | 544 | 37.0 |
| 5k-7k | MP | 89477188 | 8947718800 | 88619814 | 630 | 37.0 |
| NoGel* | MP | 95838352 | 9583835200 | 92398224 | 536 | 37.0 |

**Supplementary Table 2**: Long-read data summary

| **Library** | **#Cells** | **#reads (>1000 bp)** | **N50** | **Total GBs** |
| --- | --- | --- | --- | --- |
| PacBio RS II | 18 | 1948454 | 10456 | 16450583878 |
| Minion R7.4 | 6 | 450463 | 4916 | 1683754781 |

**Supplementary Table 3**: *Maniola jurtina* genome characteristics

| **Property** | **Minimum** | **Maximum** |
| --- | --- | --- |
| Heterozygosity | 1.89416% | 1.9291% |
| Genome Haploid Length | 573,649,617 bp | 576,419,021 bp |
| Genome Repeat Length | 433,786,156 bp | 435,880,342 bp |
| Genome Unique Length | 139,863,460 bp | 140,538,678 bp |
| Model Fit | 96.0281% | 98.7053% |
| Read Error Rate | 0.263087% | 0.263087% |

**Supplementary Table 4**: *M. jurtina* genome properties

| **Properties** | **Genome** |
| --- | --- |
| # contigs (> 0 bp) | 20143 |
| # contigs (> 1000 bp) | 10860 |
| # contigs (> 5000 bp) | 7206 |
| # contigs (>= 10000 bp) | 5836 |
| # contigs (>= 25000 bp) | 4345 |
| # contigs (>= 50000 bp) | 2744 |
| Total length (>= 0 bp) | 623665947 |
| Total length (>= 1000 bp) | 618415580 |
| Total length (>= 5000 bp) | 611197145 |
| Total length (>= 10000 bp) | 601622982 |
| Total length (>= 25000 bp) | 576227151 |
| Total length (>= 50000 bp) | 518313756 |
| Largest contig | 2944739 |
| Total length | 618415580 |
| GC (%) | 36.90 |
| N50 | 214423 |
| N75 | 78459 |
| L50 | 658 |
| L75 | 1875 |
| # N's per 100 kbp | 8864.86 |

**Supplementary Table 5**: BUSCO summary

| **BUSCO set** | **Total BUSCO** | **Complete**  **single-copy** | **Complete** **duplicated** | **Fragmented** | **Missing** |
| --- | --- | --- | --- | --- | --- |
| Arthropoda | 1066 | 963 | 219 | 37 | 66 |
| Insecta | 1658 | 1143 | 329 | 58 | 128 |
| Eukaryota | 303 | 198 | 71 | 14 | 20 |

**Supplementary Table 6**: CEGMA summary

| **CEGMA (248)** | **#Prots** | **%Completeness** | **#Total** | **Average** | **%Ortho** |
| --- | --- | --- | --- | --- | --- |
| Complete | 198 | 79.84 | 339 | 1.71 | 46.46 |
| Partial | 227 | 91.53 | 479 | 2.11 | 65.20 |

**Supplementary Table 7**: *M. jurtina* genome annotation summary

| **Properties** | **Value** |
| --- | --- |
| Number of genes | 36,226 |
| Number of mrnas | 38,067 |
| Number of mrnas with utr both sides | 404 |
| Number of mrnas with at least one utr | 2,663 |
| Number of cdss | 38,067 |
| Number of exons | 177,284 |
| Number of five_prime_utrs | 1,799 |
| Number of introns | 142,284 |
| Number of start_codons | 36,008 |
| Number of stop_codons | 36,485 |
| Number of three_prime_utrs | 1,268 |
| Number of exon in cds | 177,284 |
| Number of exon in five_prime_utr | 1,799 |
| Number of exon in three_prime_utr | 1,268 |
| Number of intron in cds | 139,217 |
| Number of intron in exon | 139,217 |
| Number of intron in intron | 111,757 |
| Number of single exon gene | 8,433 |
| Number of single exon mrna | 8,509 |
| mean mrnas per gene | 1.10 |
| mean cdss per mrna | 1.00 |
| mean exons per mrna | 4.70 |
| mean five_prime_utrs per mrna | 0.00 |
| mean introns per mrna | 3.70 |
| mean start_codons per mrna | 0.90 |
| mean stop_codons per mrna | 1.00 |
| mean three_prime_utrs per mrna | 0.00 |
| mean exons per cds | 4.70 |
| mean exons per five_prime_utr | 1.00 |
| mean exons per three_prime_utr | 1.00 |
| mean introns in cdss per mrna | 3.70 |
| mean introns in exons per mrna | 3.70 |
| mean introns in introns per mrna | 2.90 |
| Total gene length | 150,339,799 |
| Total mrna length | 167,875,458 |
| Total cds length | 37,192,228 |
| Total exon length | 39,884,925 |
| Total five_prime_utr length | 1,833,292 |
| Total intron length | 130,683,230 |
| Total start_codon length | 108,024 |
| Total stop_codon length | 109,455 |
| Total three_prime_utr length | 859,405 |
| Total intron length per cds | 128,129,750 |
| Total intron length per exon | 128,129,750 |
| Total intron length per intron | 20,463,359 |
| mean gene length | 4,150 |
| mean mrna length | 4,409 |
| mean cds length | 977 |
| mean exon length | 224 |
| mean five_prime_utr length | 1,019 |
| mean intron length | 918 |
| mean start_codon length | 3 |
| mean stop_codon length | 3 |
| mean three_prime_utr length | 677 |
| mean cds piece length | 209 |
| mean five_prime_utr piece length | 1,019 |
| mean three_prime_utr piece length | 677 |
| mean intron in cds length | 920 |
| mean intron in exon length | 920 |
| mean intron in intron length | 183 |
| Longest genes | 93,735 |
| Longest mrnas | 93,735 |
| Longest cdss | 57,939 |
| Longest exons | 11,979 |
| Longest five_prime_utrs | 8,414 |
| Longest introns | 36,945 |
| Longest start_codons | 3 |
| Longest stop_codons | 3 |
| Longest three_prime_utrs | 5,878 |
| Longest cds piece | 11,979 |
| Longest five_prime_utr piece | 8,414 |
| Longest three_prime_utr piece | 5,878 |
| Longest intron into cds part | 36,946 |
| Longest intron into exon part | 36,946 |
| Longest intron into intron part | 11,980 |
| Shortest genes | 152 |
| Shortest mrnas | 152 |
| Shortest cdss | 40 |
| Shortest exons | 3 |
| Shortest five_prime_utrs | 35 |
| Shortest introns | 6 |
| Shortest start_codons | 3 |
| Shortest stop_codons | 3 |
| Shortest three_prime_utrs | 6 |
| Shortest cds piece | 3 |
| Shortest five_prime_utr piece | 35 |
| Shortest three_prime_utr piece | 6 |
| Shortest intron into cds part | 43 |
| Shortest intron into exon part | 43 |
| Shortest intron into intron part | 7 |

**Supplementary Table 8:** OrthoFinder Summary

| **Property** | **Value** |
| --- | --- |
| Number of genes | 349442 |
| Number of genes in orthogroups | 302123 |
| Number of unassigned genes | 47319 |
| Percentage of genes in orthogroups | 86.5 |
| Percentage of unassigned genes | 13.5 |
| Number of orthogroups | 15064 |
| Number of species-specific orthogroups | 670 |
| Number of genes in species-specific orthogroups | 2967 |
| Percentage of genes in species-specific orthogroups | 0.8 |
| Mean orthogroup size | 20.1 |
| Median orthogroup size | 18 |
| G50 (assigned genes) | 26 |
| G50 (all genes) | 23 |
| O50 (assigned genes) | 3476 |
| O50 (all genes) | 4439 |
| Number of orthogroups with all species present | 2915 |
| Number of single-copy orthogroups | 39 |

**Supplementary Table 9:** Divergence time of major nodes used as fossil calibration in this study.

| **Groups** | **Approx. divergence time (per 100 MY)** | **Reference** |
| --- | --- | --- |
| Trichoptera - Lepidoptera | **B(2.75, 3.0)** | Tong et al. Science 349 (6247), 487 (2015) |
| *P. xylostella* – *B. mori* | **B(1.40, 1.50)** | Tong et al. Science 349 (6247), 487 (2015) |
| *O. brumata* – *P. polytes* | **B(1.15, 1.25)** | Tong et al. Science 349 (6247), 487 (2015) |
| *B. mori* – *P. interpunctella* | **B(1.10, 1.15)** | Tong et al. Science 349 (6247), 487 (2015) |
| *C. cecrops* – *D. plexippus* | **B(0.85, 0.95)** | Tong et al. Science 349 (6247), 487 (2015) |
| *P. polytes* - Nymphalidae | **B(1.0, 1.10)** | Tong et al. Science 349 (6247), 487 (2015) |
| Diptera – Lepidoptera - Trichoptera | **@3.5** | Tong et al. Science 349 (6247), 487 (2015) |
